# Limited Expression of *Nrf2* in Neurons Across the Central Nervous System

**DOI:** 10.1101/2023.05.09.540014

**Authors:** Daniel C. Levings, Salil Saurav Pathak, Yi-Mei Yang, Matthew Slattery

## Abstract

Nrf2 is a broadly expressed transcription factor that regulates gene expression in response to reactive oxygen species (ROS) and oxidative stress. It is commonly referred to as a ubiquitous pathway, but this generalization overlooks work indicating that Nrf2 is essentially unexpressed in some neuronal populations. To explore whether this pattern extends throughout the central nervous system (CNS), we quantified *Nrf2* expression and chromatin accessibility at the *Nrf2* locus across multiple single cell datasets. In both the mouse and human CNS, *Nrf2* was repressed in almost all mature neurons, but highly expressed in non-neuronal support cells, and this pattern was robust across multiple human CNS diseases. A subset of key Nrf2 target genes, like *Slc7a11*, also remained low in neurons. Thus, these data suggest that while most cells express Nrf2, with activity determined by ROS levels, neurons actively avoid Nrf2 activity by keeping *Nrf2* expression low.

## Introduction

Oxidative metabolism is inextricably linked to the production of reactive oxygen species (ROS), which have the potential to damage all classes of macromolecules [1-4]. Cellular ROS are byproducts of energetic pathways, but they are also generated upon exposure to many xenobiotics. Excess ROS generate oxidative stress and promote diseases ranging from cancer to neurodegeneration, so ROS are often viewed as cytotoxic [1-4]. Yet ROS are not invariably detrimental. Several properties make ROS useful signaling molecules, including their potential for rapid modification of proteins and close ties to cellular metabolism, and they have been co-opted as signaling molecules in many cellular processes, including stem cell self-renewal and differentiation [5-10]. Thus, ROS are unavoidable, necessary, and potentially dangerous, so maintaining balanced ROS levels is essential. Accordingly, cells have antioxidant pathways, like the Nrf2 pathway in metazoans, that keep ROS at cell type-appropriate levels and mitigate their damaging effects [11].

Nrf2, encoded by the gene *Nfe2l2* (*Nuclear factor erythroid-derived 2-like 2*), is a widely expressed transcription factor that is often referred to as the ‘master regulator’ of the response to oxidative stress [12-16]. When ROS levels are low, Nrf2 is largely sequestered in the cytoplasm by the inhibitory protein Keap1, which binds Nrf2 and targets it for proteasomal degradation [17-20]. Keap1 is sensitive to ROS, however, which modify several reactive cysteines on Keap1 and impair its ability to bind Nrf2. As ROS levels increase, Keap1-mediated Nrf2 inhibition decreases, and nuclear Nrf2 increases. In the nucleus, Nrf2 dimerizes with one of the small Maf (v-maf avian musculoaponeurotic fibrosarcoma oncogene homolog) proteins, binds to *cis*-regulatory antioxidant response elements (AREs), and drives upregulation of a range of target genes [20, 21]. Nrf2 is known for its regulation of antioxidant genes, but it also regulates genes involved in drug/xenobiotic metabolism, carbohydrate metabolism, and more [12]. Activation of Nrf2 is cytoprotective and promotes longevity, while loss of Nrf2 leads to increased sensitivity to drivers of cancer and neurodegeneration, and decreased lifespan [22-27].

Like most stress-responsive transcription factors, Nrf2 is broadly expressed. It is often referred to as having ubiquitous activity [28-36]. However, although most recognize that Nrf2 activity varies across tissues, the ‘ubiquitous’ designation does not fit with important work indicating that Nrf2 is essentially unexpressed and inactive in some neuronal populations. A dearth of Nrf2 activity in neurons was first implied by the lack of ARE *cis*-regulatory activity in cultured mouse cerebellar granule cells and cortical neurons [37-40]. Further work demonstrated that *Nrf2* (transcript and protein) is dramatically repressed in cultured rodent cortical neurons when compared to astrocytes [41-43]. It is not yet clear whether this *Nrf2* expression pattern extends far beyond cortical neurons to most other neural cell types. But considering Nrf2’s central role in counteracting oxidative stress, which is a key driver of multiple neurodegenerative diseases, it is important to understand whether its neuronal repression is generalizable across the central nervous system (CNS). Here we used multiple single cell genomic datasets to explore *Nrf2* expression and regulation in hundreds of neuronal and non-neuronal cell types in mouse and human. With few exceptions, *Nrf2* is expressed at far lower levels in neurons than in non-neuronal support cells in both species, this pattern is maintained in multiple disease states, and the chromatin accessibility landscape at the *Nrf2* locus parallels these expression differences. These results imply that Nrf2 activity is limited in almost all neurons of the mouse and human CNS.

## Results

### Low expression of Nrf2 in almost all neurons of the CNS

To gain a quantitative view of *Nrf2* expression in all cell types across the CNS, we used single cell RNA sequencing (scRNA-seq) data from mice [44-46] and humans [47]. Because of the sparsity of scRNA-seq data, for each dataset we analyzed aggregate data where sequencing reads from all individual cells of a given cell type are combined to generate a higher depth transcriptional profile of that cell type. We then separated cell types into neuron or non-neuronal “support” cell categories; the general “support” term is not meant to minimize the functional relevance of non-neuronal cells in the CNS, but is an umbrella term meant to cover everything from glial cell types (astrocytes, microglia, oligodendrocytes) to endothelial cells. In the scRNA-seq dataset from adolescent mice [45], *Nrf2* expression was significantly lower overall in neurons as compared to non-neuronal support cells **(Figure 1A)**. A similar pattern, with little-to-no *Nrf2* expression in CNS neurons, was observed in an independent adult mouse dataset **(Figure S1)** [44].

**Figure 1.**
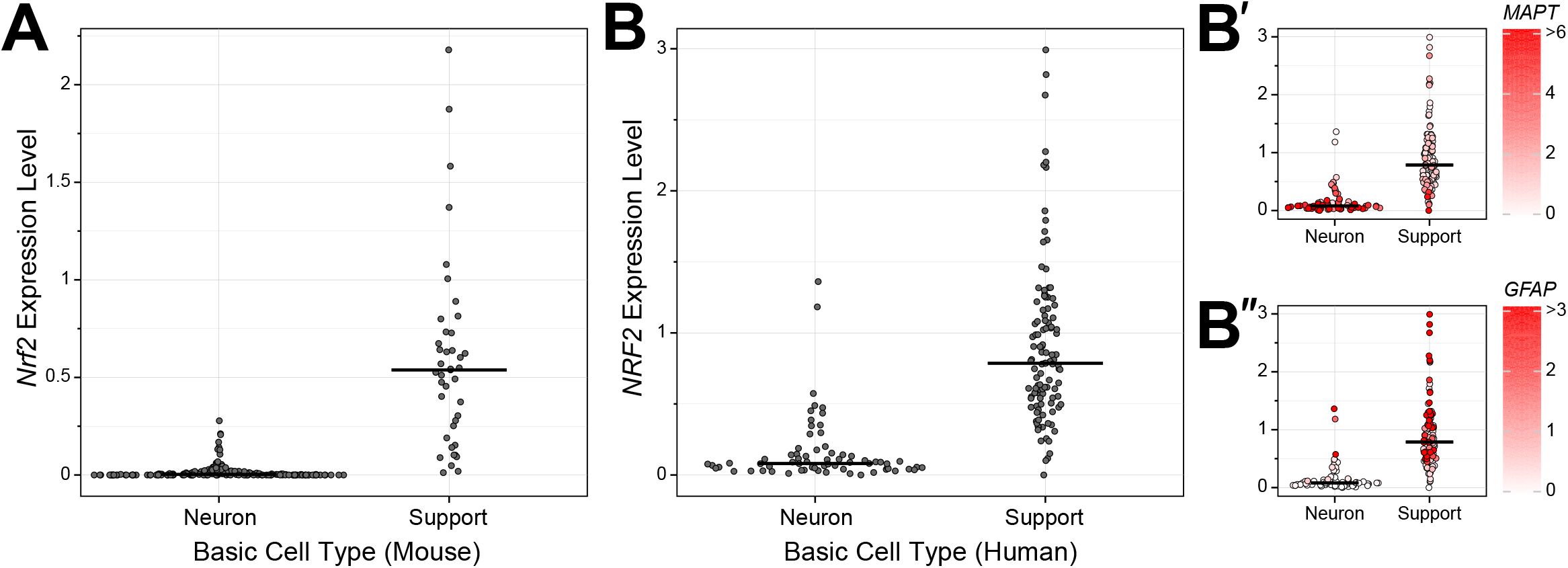
*Nrf2* expression in neurons versus support cells. (A) Dot plot summarizing aggregate scRNA-seq analysis of *Nrf2* expression in the adolescent mouse brain. All sequencing reads from individual cells of a given cell type were combined, and each dot represents Nrf2 levels in a cell type (see Methods). Cell types are separated based on classification as neurons or support cells, and median expression for each class is indicated with a black horizontal line. Data are from [45]. (B) Same as (A), only with aggregate analysis of *NRF2* expression in the human brain. Expression levels of the neuronal marker *MAPT* **(B’)** and astrocyte marker *GFAP* **(B’’)**, calculated using the same approach used for *NRF2*, are overlaid on each cell type dot in the panels on the right. Data are from [47].

We next used human scRNA-seq data [47] to ascertain if widespread downregulation of *NRF2* in neurons also occurs in the human brain. Like the pattern observed in mice, *NRF2* was significantly downregulated in human neurons **(Figure 1B)**. However, in this case there were multiple cell types originally annotated as neurons that express *NRF2* at levels comparable to support cells. Further exploration of these ‘high *NRF2*’ neurons revealed that, unlike most other neurons, they expressed low levels of the neuronal markers *MAPT* and *RBFOX2*, and high levels of the glial markers *GFAP, AQP4*, and *SLC1A3* **(Figure 1B and S2A-C)**. Therefore, although these ‘high *NRF2*’ cells were annotated as neurons in the original dataset, they do not express key neural marker genes and do express several important glial marker genes. Consistent with this, these cell populations did not cluster with neurons in whole transcriptome-based principal component analysis **(Figure S2D)**, suggesting these cells may be more accurately annotated as non-neuronal or neural precursor cell types. To determine if *NRF2*’s expression pattern is maintained in pathological states, we analyzed brain single nucleus (sn)RNA-seq data from patients with Alzheimer’s disease [48], multiple sclerosis [49], or autism [50] – three neurological disorders associated with oxidative stress in the brain [51-55]. The *NRF2* expression differences in neuron versus support cells were clear in all three disease states **(Figure 2)**. Ultimately, the human *NRF2* expression pattern is similar to that seen in the mouse brain, with *NRF2* unexpressed in most neurons and expressed in most support cells, and this pattern is maintained across multiple pathologies.

**Figure 2.**
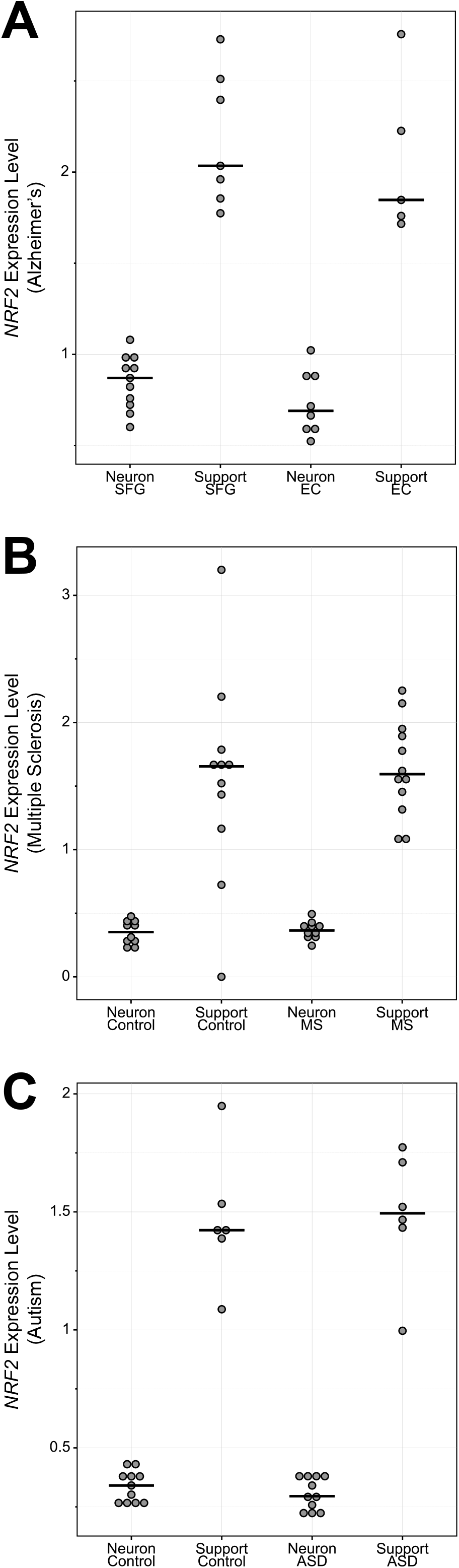
*NRF2* expression in neurons versus support cells in human neurological disorders. (A) Dot plot summarizing aggregate snRNA-seq data from nuclei isolated from postmortem brains of individuals with Alzheimer’s disease [48]. Nuclei were isolated separately from the superior frontal gyrus (SFG) or entorhinal cortex (EC). As in previous figures, cell types are broken down by classification as neurons or support cells. (B) Similar to (A), only with snRNA-seq data from postmortem multiple sclerosis (MS) brain tissue samples and controls [49]. Nuclei were isolated from the cortical gray matter and subcortical white matter MS lesion areas. (C) Similar to (A), only with snRNA-seq data from postmortem autism spectrum disorder (ASD) brain tissue samples and controls [50]. Nuclei were isolated from the prefrontal cortex and anterior cingulate cortex.

To understand when in the differentiation process Nrf2 is downregulated, we used an embryonic mouse brain dataset that allowed for further resolution of *Nrf2* expression across stages of neuronal differentiation [46]. Consistent with the above results, *Nrf2* expression was markedly downregulated in CNS neurons (relative to support cells) in the embryonic mouse brain **(Figure 3**; log10 scale in **Figure S3)**. It was expressed highly in embryonic precursor cells of the neural crest/tube, moderately in embryonic neuroblasts, and minimally in embryonic neurons. Further, for differentiated neurons, *Nrf2* expression was higher in early embryogenesis and decreased as development progressed (not shown). Thus, throughout brain development, *Nrf2* was expressed in most support cells and neural precursors, and virtually unexpressed in neurons.

**Figure 3.**
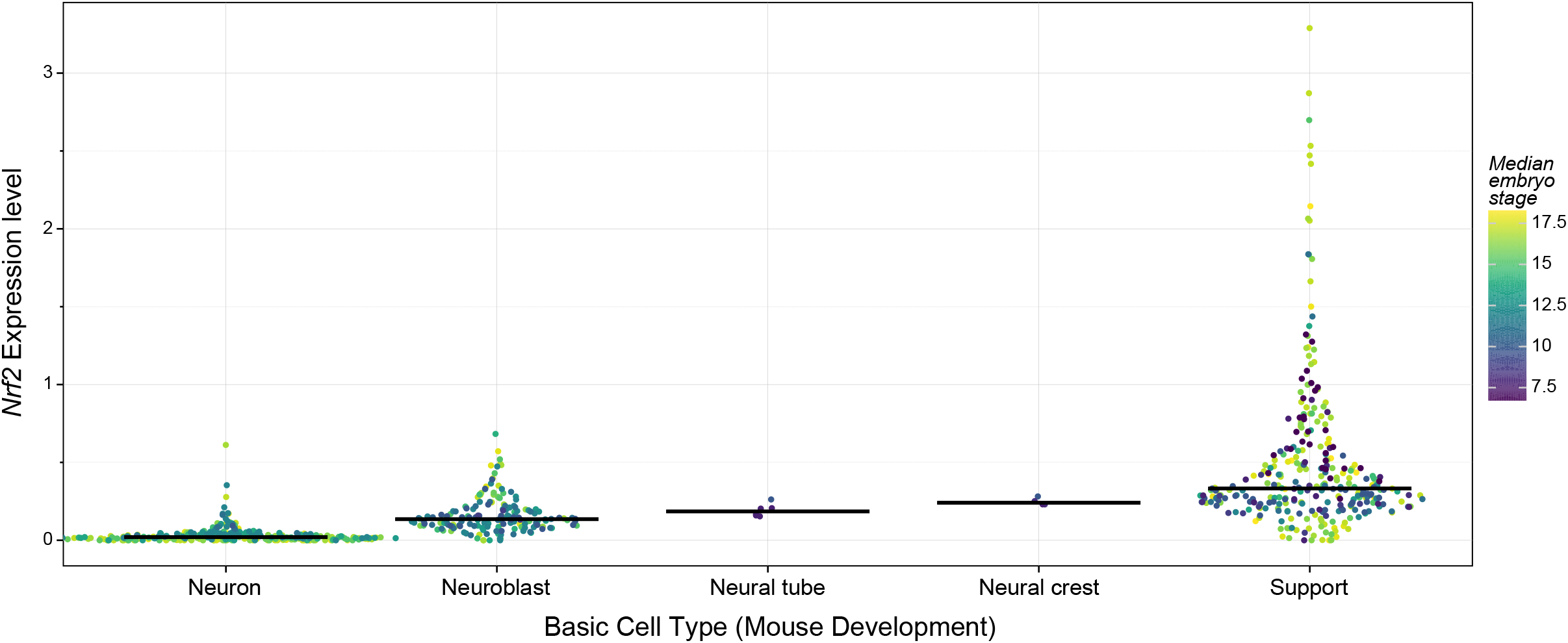
*Nrf2* expression in neurons versus support cells across mouse development. Dot plot summarizing aggregate scRNA-seq analysis of *Nrf2* expression in the developing mouse brain. Cell types are separated based on their classification as neurons, neuroblasts, neural tube cells, neural crest cells, and support cells as indicated. Dot color represents the median embryonic stage (e7 through e18) for the population of cells assigned to each cell type. The same graph is presented with expression on a log scale in Figure S4. Data are from [46].

### Repressive chromatin at the Nrf2 locus in CNS neurons

The above data indicate that the mechanism(s) maintaining low NRF2 in neurons are robust and conserved in mammals. We explored whether the differences might be maintained at the chromatin level using ATAC-seq (Assay for Transposase-Accessible Chromatin using sequencing) data, which allows for genome-wide monitoring of chromatin accessibility [56]. ATAC-seq data can complement RNA-seq data because in order for a gene to be expressed, its promoter must be accessible to the general transcription machinery. We used aggregated single nucleus (sn)ATAC-seq data from the mouse brain [57], to determine if the *Nrf2* promoter or any other regulatory DNA regions at its locus differ in accessibility between neurons and support cells. Indeed, the *Nrf2* promoter and an intronic enhancer just downstream of the promoter were highly accessible in support cells and inaccessible across almost all types of neurons **(Figure 4** and **S4)**. The differences were dramatic enough that unbiased clustering of the accessibility profiles across the *Nrf2* locus was able to clearly separate support cells from neuronal cell types **(Figure S5)**, with the exception of immature oligodendrocytes that clustered with neurons. Mature oligodendrocytes, however, clustered with support cells. Overall, these snATAC-seq data indicate that in the CNS neurons have a repressive chromatin accessibility landscape at *Nrf2*, whereas the landscape in non-neuronal cells is permissive for *Nrf2* expression.

**Figure 4.**
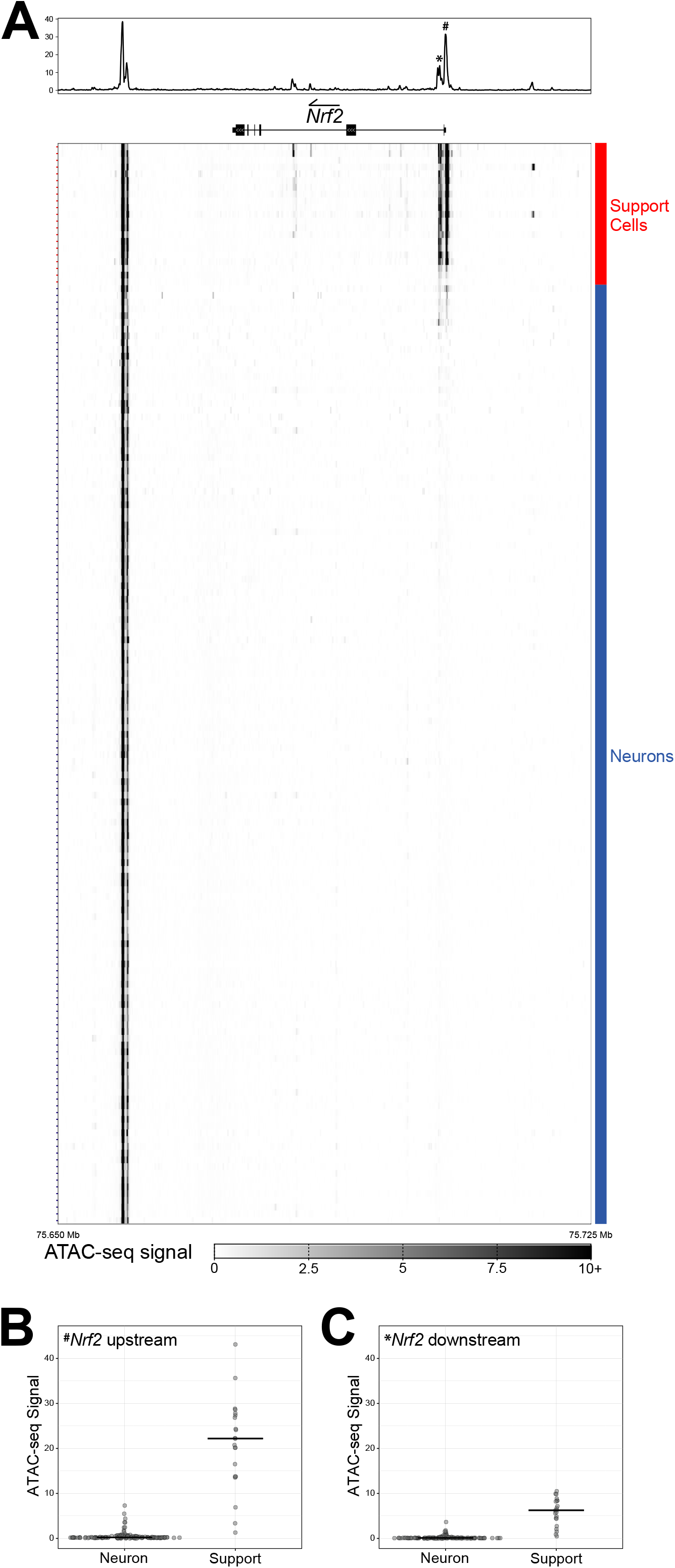
*Nrf2* locus chromatin environment in neurons versus support cells. (A) Heatmap of the aggregate snATAC-seq signal at the *Nrf2* (i.e., *Nfe2l2*) locus across cell types of the mouse brain. As with the scRNA-seq data, sequencing reads from individual cells of a given cell type were combined to generate a complete profile for each cell type. The cumulative read profile is represented at the top, above an aligned model of the *Nrf2* gene (arrow indicates the direction of Nrf2 coding). The pound sign (#) in the cumulative read profile marks the prominent peak upstream of Nrf2 transcription start site, and asterisk (*) marks the downstream peak in Nrf2’s first intron – both are addressed in panels (B) and (C). Rows of the heatmap represent the aggregate snATAC-seq signal of each cell type, which are classified as neurons (blue) or support cells (red) as indicated to the right of the heatmap. The genomic region shown is zoomed out to include a positive control region downstream of *Nrf2* at the promoter of *Hnrnpa3* (not shown in gene model) that is highly accessible across all cell types. Data are from [57]. (B) Dot plot representing the aggregate snATAC-seq signal across cell types for the accessible region upstream (#) of the *Nrf2* transcription start site. Cell types are separated based on classification as neurons or support cells, and values are based on the maximum ATAC-seq signal for this peak region. (C) Same as (B), only with dot plot representing the aggregate snATAC-seq signal across cell types for the accessible region downstream (*) of the *Nrf2* transcription start site.

## Discussion

Previous work suggested that the ROS-responsive transcription factor *Nrf2* is not expressed in subsets of mouse neurons [37-43]. To explore whether this pattern extends throughout the brain, we quantified (1) *Nrf2* expression across multiple CNS scRNA-seq and snRNA-seq datasets from mice and humans, and (2) chromatin accessibility at the Nrf2 locus in independent CNS snATAC-seq data. Ultimately, we found that this pattern is generalizable throughout the mammalian CNS: *Nrf2* is repressed in almost all mature neurons, but expressed in most non-neuronal support cells.

There is a clear developmental component to *Nrf2* repression, as it is expressed in embryonic precursor cells of the neural crest/tube and neuroblasts, and in cells undergoing adult neurogenesis (not shown), which is consistent with its potential function in neural progenitor cells [32]. And its repression as differentiation progresses is important because forced expression of Nrf2 in cortical neurons *in vitro* hinders normal development of these neurons [42]. The ATAC-seq data, which are in line with previous work suggesting this repression is mediated at the chromatin level [42], suggest that putative regulatory DNA regions in *Nrf2*’s promoter and first intron are important for the differential expression between support cells and neurons. Importantly, Nrf2 positively autoregulates via its promoter region [59], and we previously found that the accessible region in intron 1 is also bound and positively regulated by NRF2 [58]. Thus, it is likely that disruption of this Nrf2 autoregulatory loop plays a role in its downregulation in neurons. One possible disruptor of this loop is the Nrf2 inhibitor Keap1 and, consistent with this, *Keap1* increases as *Nrf2* decreases with neuronal differentiation **(Figure S6)**.

Regardless of the mechanism, it is not clear why an important, near ubiquitous cytoprotective transcription factor like Nrf2 remains off in mature neurons. Especially considering oxidative stress is a driver of many diseases [52, 60]. The simplest explanation is that Nrf2 activity also disrupts the normal function of mature neurons. Notably, ROS play a key role in controlling synaptic plasticity in mature neurons [61-65]. And these activity-dependent changes in synaptic transmission, which are important for learning and memory, are disrupted by antioxidants [66]. A subset of important Nrf2-targeted antioxidant genes (e.g., *Slc3a2, Slc7a11, Nqo1, Prdx1*) are also low in neurons **(Figure S7)**, so it is likely that these and/or other Nrf2 targets must remain low or non-ROS-responsive in mature neurons.

Stress response pathways are often viewed as ubiquitous, with activity determined primarily by stress conditions. However, it is becoming increasingly clear that stress responses should be tailored to fit the precise needs of a particular cell type [67]. In the case of Nrf2, this tailoring has resulted in its specific repression in neurons. Future work exploring *why* this expression pattern persists in mature neurons will inform our models on the roles of antioxidant genes in normal neuronal physiology and in neurological disorders.

## Materials and Methods

### Expression comparisons of neural and support cell types

For all figures comparing expression levels of support cells and neural cell types, the average expression in raw counts for every gene per cell type was either previously reported (Mouse Brain Atlas [45] and DropViz [44] data) or calculated ad-hoc for each dataset – see “Aggregation of single cell expression by cell type from human data” section below for a detailed description of how these were generated. These values were imported into Python (version 3.9.13) using either the *loompy* package (version 3.0.7, https://github.com/linnarsson-lab/loompy) or as text files, and were library-size normalized by dividing the average raw counts for each gene by the total of the average gene counts across all genes in a given cell type (i.e., dividing each element by the column sum). These average library size-normalized expression values were then multiplied by 10000 in order to plot expression on similar scales across all single cell RNA-seq datasets.

To plot the expression differences of the figures listed above, the cell metadata was imported as a ‘plotting data frame’ and each cell type was annotated as a ‘neural’ or ‘support’ cell using column data in the metadata files in combination with previously published information concerning these neural/other cell types in the relevant datasets. The appropriate row(s) of gene expression values were then isolated from the library size-normalized gene-by-cell-type expression matrix and added to this ‘plotting data frame’. For the mouse developmental data from the Linnarsson lab’s Mouse Brain Atlas developmental dataset [46], the counts for cells of each embryonic stage (e7.0 to e18.0) were used to calculate a median embryonic stage for each cell type in the dataset (Figures 3, S3 and S6). The *agg()* function of the *pandas* package [68] was used to summarize expression levels for neural/support cells, or other relevant grouping, by median expression level. Finally, the *plotnine* python package [69] was used to plot the normalized gene expression values for each cell type within each group on the x-axis using Sina plots [70], *geom_sina()* function, and the *geom_errorbar()* function for displaying median expression.

### DropViz expression comparison

Cell type-aggregated expression levels for *NRF2* were obtained from DropViz (http://dropviz.org) for mouse brain single cell RNA-seq using the following steps: 1) on the ‘Query’ tab, *Nfe2L2* was entered as Gene and ‘Update!’ was selected, 2) the ‘Global Clusters’ tab was opened as a ‘Table’, and 3) the ‘Download as CSV’ button was used to download a comma-separated table (CSV file) of expression values for *Nfe2l2* by region and cell type/cluster. The expression values downloaded from DropViz and plotted here are the mean single-cell UMI gene counts per 100,000 cell type UMI counts, log10-scaled. Due to the relatively smaller number of cell types plotted (compared to the other single cell datasets), these data were plotted in Python (version 3.9.13) using dot plots, *geom_dotplot()* function, rather than sina plots but were otherwise plotted using the same methodology as the other expression comparisons.

### Aggregation of single cell expression by cell type from human data

Single cell RNA-seq data were downloaded from either UCSC Cell Browser or CZ CELLxGENE and processed using the Seurat package from *R* version 4.1.0. Data from UCSC Cell Browser included: adult human brain (https://cells.ucsc.edu/?ds=dev-brain-regions+wholebrain), multiple sclerosis (https://cells.ucsc.edu/?ds=ms), and autism (https://autism.cells.ucsc.edu) datasets. For the UCSC Cell Browser datasets, the cell metadata (*meta*.*tsv*) were loaded as data frames and the single cell expression count matrices (*exprMatrix*.*tsv*.*gz*) were loaded as sparseMatrix objects using either *fread(input = “exprMatrix*.*tsv*.*gz”)* followed by *as(x, “dgCMatrix”)* or using *ReadMtx(mtx = “rawMatrix*.*mtx*.*gz”, features = “genes*.*tsv*.*gz”, cells = “barcodes*.*tsv*.*gz”)*. The Seurat objects were then reconstituted using the *CreateSeuratObject()* function on these two pieces of data (cell metadata and count matrix). An example of this workflow is shown by UCSC Cell Browser at https://cellbrowser.readthedocs.io/en/master/load.html#seurat. Data for cells from the superior frontal gyrus (SFG) and entorhinal cortex (EC) of Alzheimer’s disease patients were downloaded as Seurat RDS objects directly from CZ CELLxGENE (https://cellxgene.cziscience.com/collections/180bff9c-c8a5-4539-b13b-ddbc00d643e6). With the Seurat objects for each of these datasets, the average expression (from raw counts) for each gene per cell type was then calculated using the *AverageExpression()* function from Seurat on each object/dataset, with slot = “counts”, and grouping on the following column/cell data: adult human brain = “cell_cluster”, multiple sclerosis = “cell_type” & “diagnosis”, autism = “cluster” & “diagnosis”, and Alzheimer’s disease = “clusterCellType”. These final gene-by-cell-type data were then exported to tab-separated tables for plotting with Python.

### Heatmap and clustering of aggregated mouse brain single nuclie ATAC-seq data

Single nuclei ATAC-seq data for mouse cerebrum were obtained from the Cis-element Atlas (CATlas, http://catlas.org/catlas_mousebrain/) [57]. Specifically, we downloaded the cell type-aggregated MACS2 signal bigWigs from CATlas (http://catlas.org/catlas_downloads/mousebrain/bigwig/). Cell type annotations and metadata were downloaded and imported from Supplementary Table 3 of Li *et al* [57] and cells were classified as support or neural in a similar fashion to previous experiments. We then extracted only the bigWig signal for a 75 Kb region including the *Nfe2l2* locus, specifically chr2: 75650000-75725000, and wrote these data to new bigWig files. These truncated bigWig files were imported into Python 3.9.13 using the *pyBigWig* package [71, 72]. Then, for each cell type-aggregated bigWig signal file, the 75 Kb signal region was broken into 2560 equal-sized bins across the entire region (in a consistent manner for all cell types) and the maximum ATAC-seq signal for each bin was computed with the *stats()* function of pyBigWig. These ‘per region bin’ max-summarized ATAC-seq signals are shown in Figures 4, S4, and S5. A subset of the max-summarized data for this 75 Kb region, including only the *Nfe2l2* gene locus and promoter region (ie-transcription start site to 5 Kb upstream of the gene), was then used for ordering cell types. For Figure 4A, the cell type annotations and max signal from an approximately ∼1.7 Kb region including the two strongest signal peaks at the *Nfe2l2* the promoter region (chr2:75703338-75705060) was used to order cell types from strongest to weakest aggregate ATAC-seq signal with the *sort_values()* function of *pandas*. For Figure S5, the bin-summarized max signal of the *Nfe2l2* locus and promoter (5 Kb upstream) only was used for ordering the cell types with the *linkage()* and *optimal_leaf_ordering()* hierarchical clustering functions of the package *scipy* [73]. The heatmaps shown in Figures 4A and S4-5 were generated using *matplotlib* [74]. Finally, the dotplots in Figure 4B-C were generated using *plotnine* and the bin-summarized max signal from the above for the promoter region: Nrf2 upstream – chr2:75704374-75705060, and Nrf2 downstream – chr2:75703338-75703975.

## Acknowledgements

The authors would like to thank Sarah Lacher and Jennifer Krznarich for helpful discussions, and Sten Linnarsson for assistance with data from the Mouse Brain Atlas. This work was supported by funding from the National Institute of General Medical Sciences (R35-GM119553 to MS) and the National Institute of Mental Health (R01-MH129300 to YMY).

## Supplementary Figure Legends

**Figure S1.**
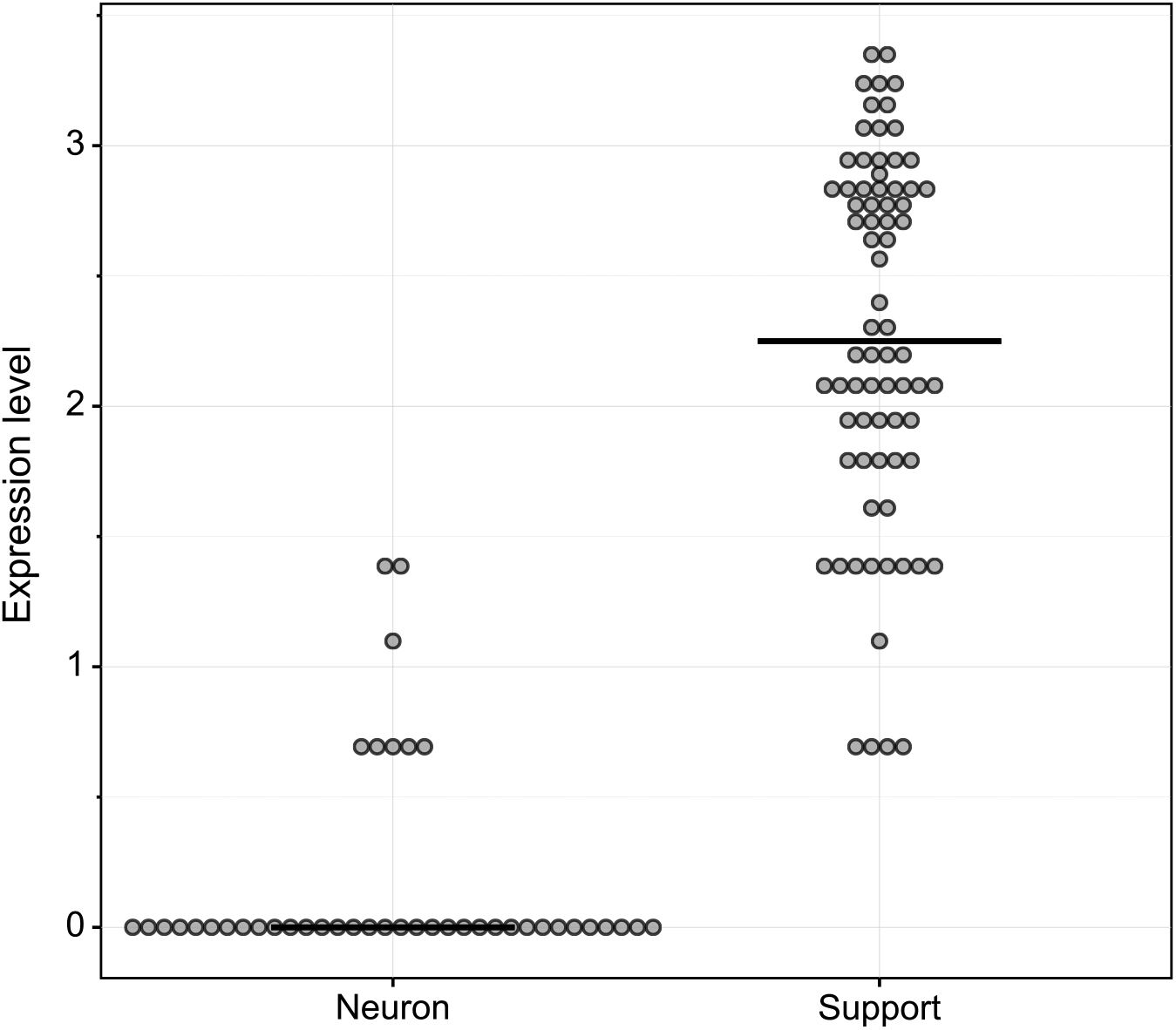
Dot plot summarizing aggregate scRNA-seq analysis of *Nrf2* expression in the adult mouse brain. As in Figure 1, cell types are separated based on classification as neurons or support cells, and median expression for each class is indicated with a black horizontal line. Data are from (Saunders et al., 2018).

**Figure S2.**
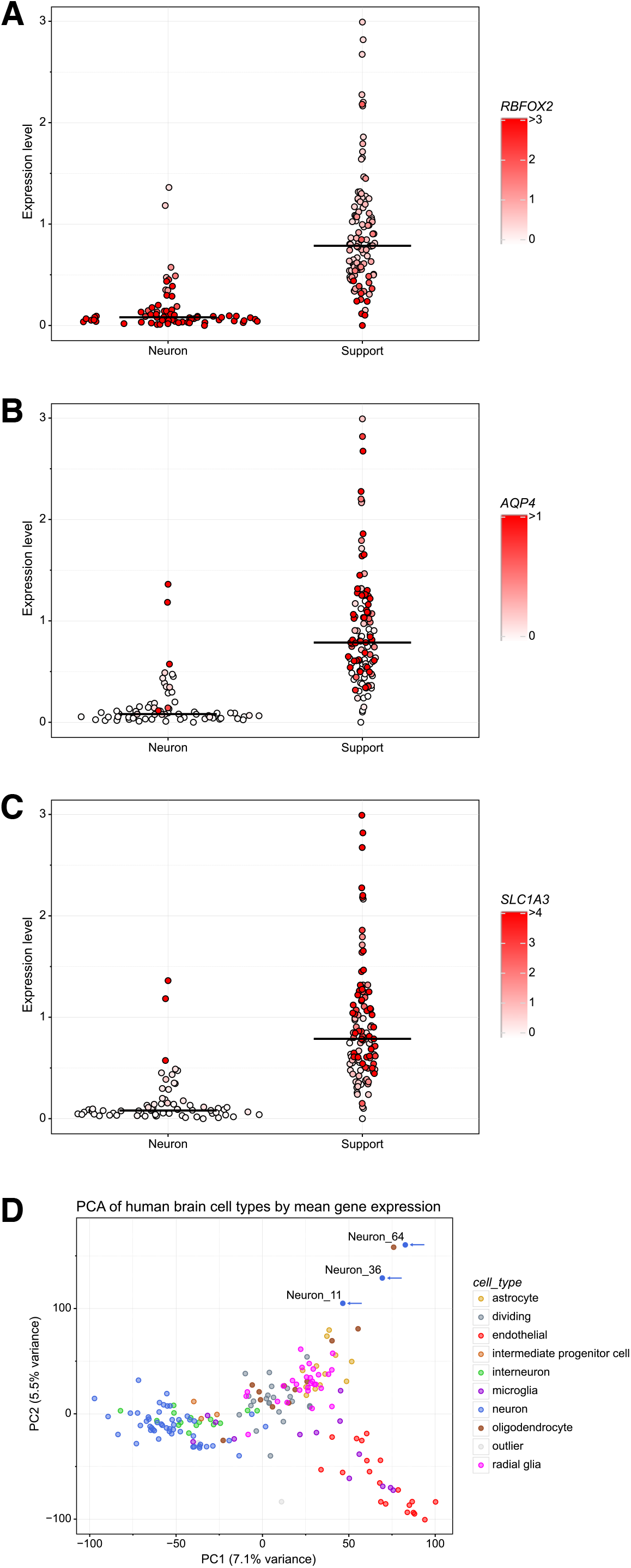
**(A - C)** Aggregate snRNA-seq analysis of *NRF2* expression in the human brain, with expression levels of **(A)** the neuronal marker *RBFOX2*, **(B)** the astrocyte marker *AQP4*, and **(C)** the astrocyte marker *SLC1A3* overlaid on each cell type dot, same as in Figure 1B. Data are from (Bhaduri et al., 2021). **(D)** Principal component analysis of aggregate cell type-summarized expression data, based on published cell type assignments (Bhaduri et al., 2021). Neuronal cell populations (blue and green dots) largely cluster separately from the other cell populations of the brain (i.e., support cells). Consistent with the patterns in panels A-C and Figure 1C, the three outlier neuron cell types with higher NRF2 expression (Neuron_11, Neuron_36, Neuron_64; marked with blue arrows) cluster separate from the other neurons and closer to support cells.

**Figure S3.**
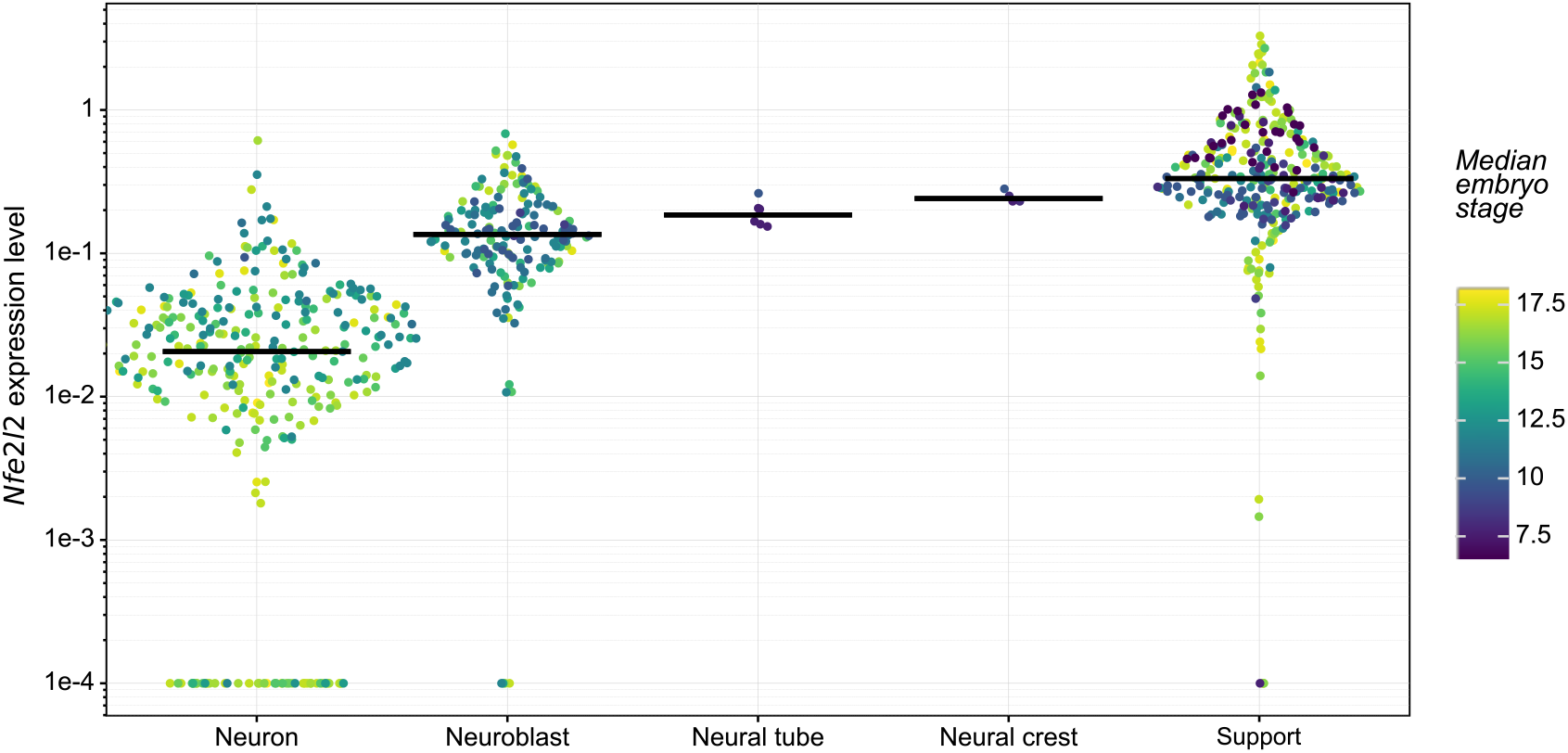
Aggregate analysis of *Nrf2* expression in the developing mouse brain. Same as Figure 1C, only with y-axis (*Nrf2* expression level) on a log_10_ scale rather than a linear scale. To display this data properly on a log_10_ scale, a pseudocount 1e-4 was added to each cell to avoid zeros.

**Figure S4.**
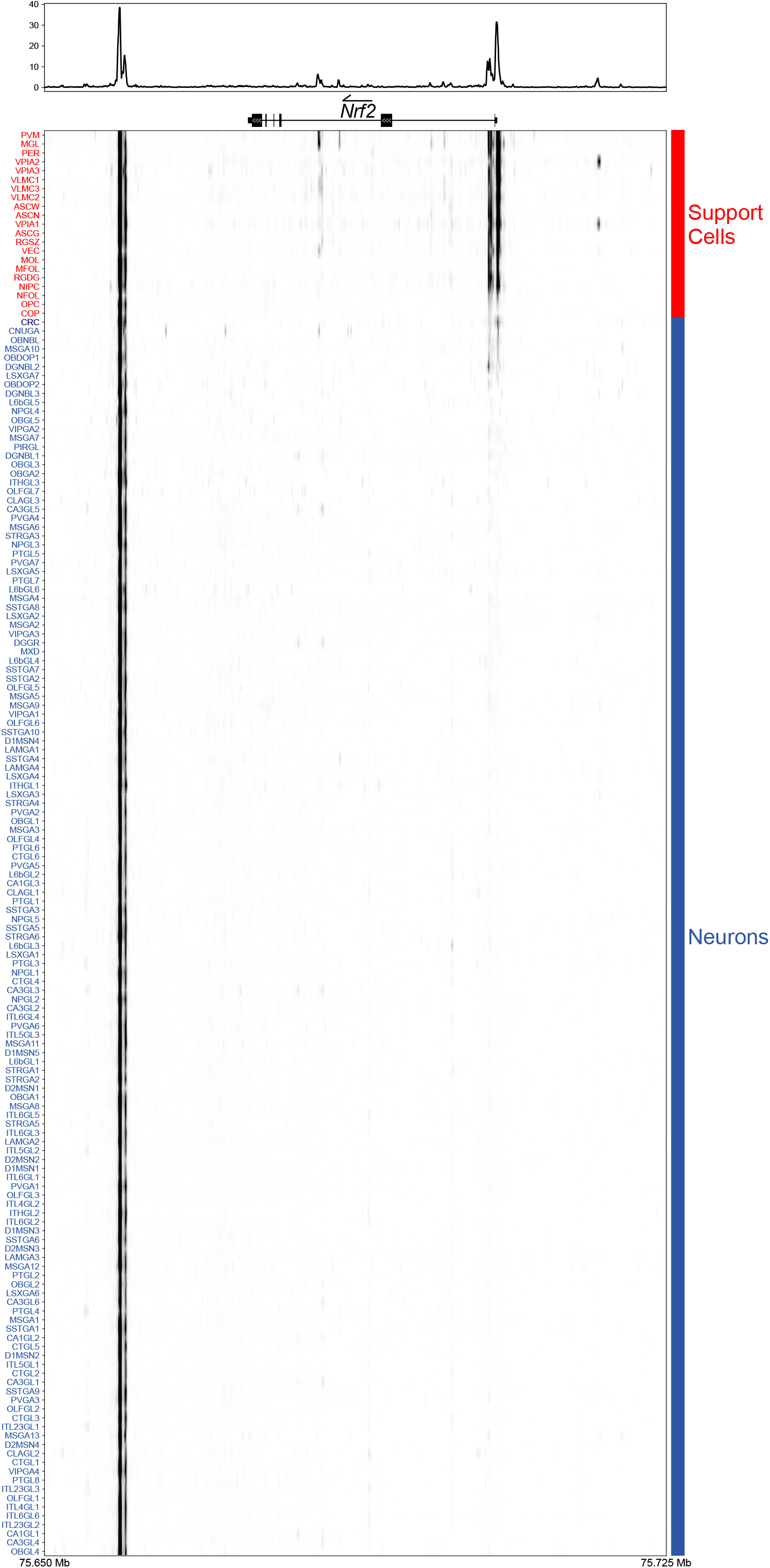
Heatmap of the aggregate snATAC-seq signal at the *Nrf2* (i.e., *Nfe2l2*) locus across cell types of the mouse brain. Same as Figure 2A, only with cell type abbreviations labeled for each row of heatmap; abbreviations are as described in (Li et al., 2021).

**Figure S5.**
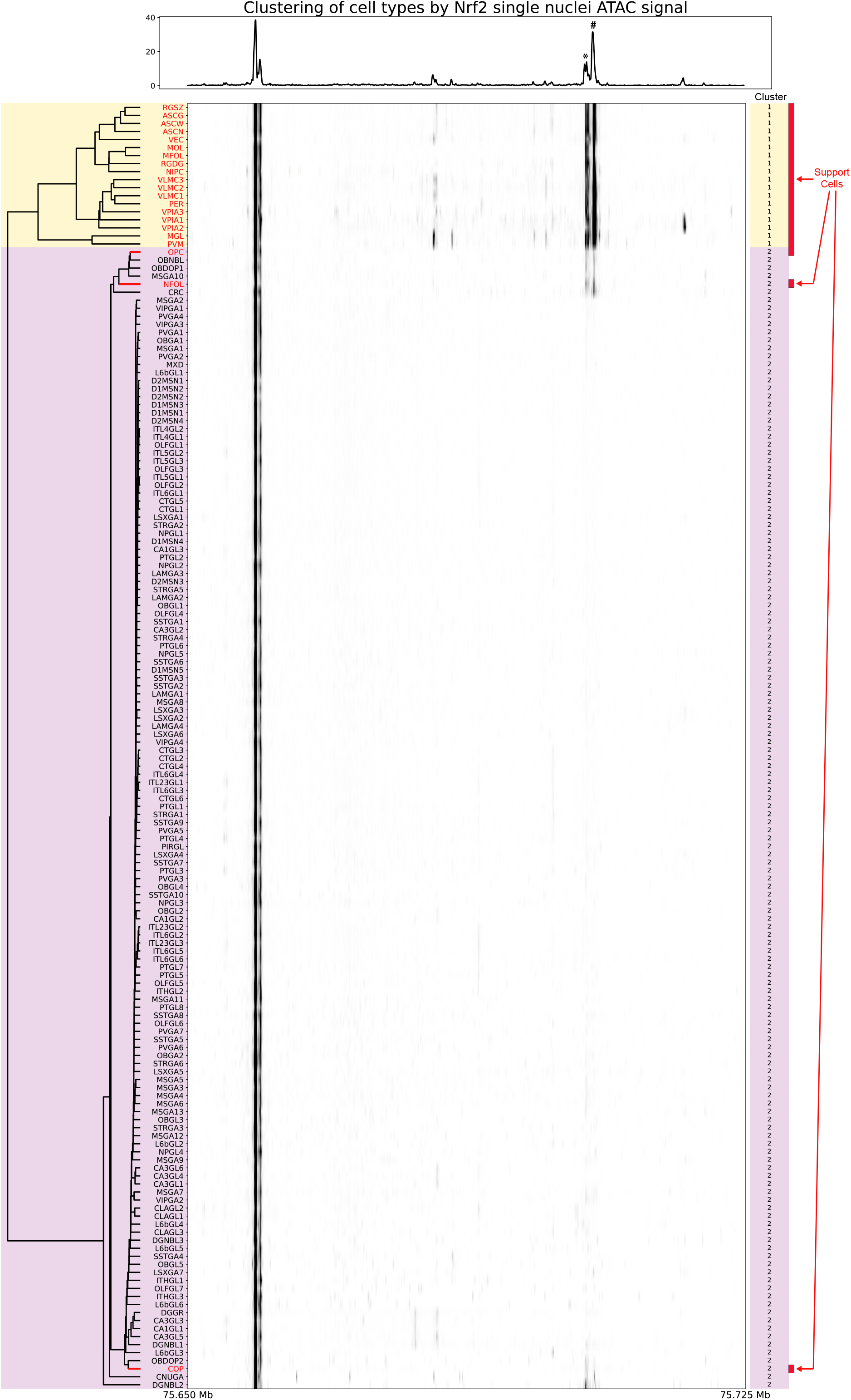
Hierarchical clustering of the aggregate snATAC-seq signal at the *Nrf2*/*Nfe2l2* gene locus. Clustering was based on the signal from the *Nrf2*/*Nfe2l2* gene body and 5 kilobases upstream of the transcription start site. The two major clusters are highlighted yellow and purple. Support cell types are indicated in red text and with red boxes to the right of the heatmap; all non-red cell types are classified as neurons.

**Figure S6.**
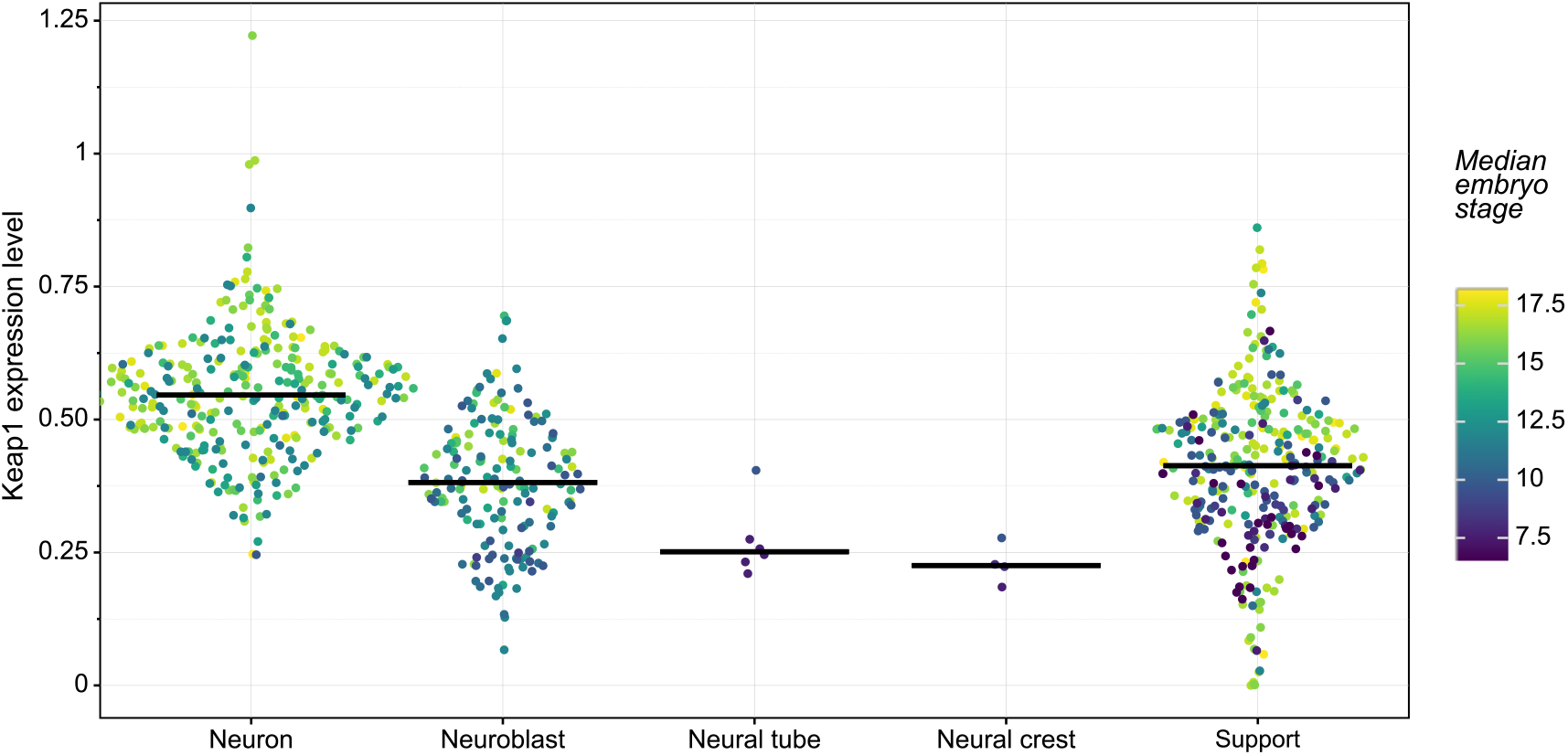
Dot plot summarizing aggregate analysis of *Keap1* expression in the developing mouse brain. As in Figure 1C, cell types are separated based on their classification as neurons, neuroblasts, neural tube cells, neural crest cells, and support cells as indicated. Dot color represents the median embryonic stage (e7 through e18) for the population of cells assigned to each cell type. Data are from (La Manno et al., 2021).

**Figure S7.**
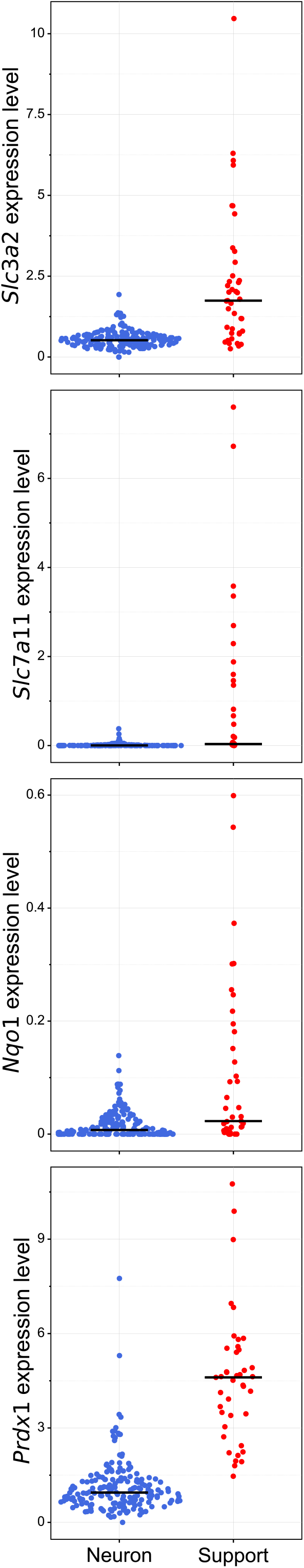
Dot plot summarizing aggregate scRNA-seq analysis of Nrf2 target gene expression in the adolescent mouse brain. Similar to Figure 1A, only four direct Nrf2 target genes (*Slc3a2, Slc7a11, Nqo1*, and *Prdx1*) are represented. Data are from (Zeisel et al., 2018).

